# Development of an Experimental Protocol to Study the Neural Control of Force and Impedance in Wrist Movements with Robotics and fMRI

**DOI:** 10.1101/2024.02.19.581013

**Authors:** Kristin Schmidt, Bastien Berret, Fabrizio Sergi

## Abstract

Robotic exoskeletons have emerged as beneficial tools in the field of rehabilitation, yet their full potential is impeded by our limited knowledge of the neural control of movements during human-robot interaction. To personalize exoskeleton protocols and improve individuals’ motor recovery, we must advance our understanding of how the brain commands movements in physical interaction tasks. However, interpreting the neural function associated with these movements is complex due to the simultaneous expression of at least two control policies: force and impedance control. This hinders our ability to isolate these control mechanisms and pinpoint their neural origins. In this study, we evaluate the capacity of externally applied forces to decouple the expression of force and impedance in a wrist-pointing task, a necessary step in isolating their neural substrates via neuroimaging.

We first conducted simulations using a neuromuscular model to examine how both force and impedance commands are updated when participants are asked to perform reaching movements in the presence of an externally applied force. Then, we recruited seven participants to perform a wrist-pointing task with the MR-SoftWrist, an MRI-compatible wrist robot. The task included four different force conditions – no force, positive constant force, negative constant force, and divergent force, each carefully selected to decouple expression of force and impedance control. Furthermore, we evaluated the efficacy of our proposed conditions for a neuroimaging experiment through simulations of neural activity. We show that these applied forces elicit distinct and predictable torque and stiffness expression, laying the groundwork for reliably identifying their associated neural activity in a future neuroimaging study.

## I. INTRODUCTION

Force and impedance control are fundamental control strategies enabling skillful human movement. Force control dominates in predictable and stable tasks, where the Central Nervous System (CNS) builds an internal model of appropriate motor commands [1]. In this case, the CNS coordinates the reciprocal activation of muscle pairs to generate desired torques. When faced with unpredictable or unstable environmental dynamics, the CNS also uses impedance control to mitigate disturbances [2], [3]. This involves the modulation of joint stiffness through the simultaneous contraction of agonist and antagonist muscle pairs. The precise regulation of force and impedance is not only essential for everyday motor tasks but also holds significant importance in expanding our knowledge of neuromuscular disorders. Following neural injuries, such as strokes, there is an alteration in the regulation of force and impedance control, often resulting in excessive co-contraction which contributes to motor impairment [4], [5]. As such, quantifying neural function responsible for specific motor control policies may lead to improved assessment of neuromuscular impairment and facilitate precision rehabilitation efforts for restoring neural function.

The role of impedance control has been well established in behavioral tasks with unstable or unpredictable dynamics [2], where the regulation of joint stiffness via co-contraction promotes task stability. However, the coactivation of agonist and antagonist muscles increases metabolic costs and can be seen as a waste of energy in tasks with predictable task dynamics. Yet, in wrist movements, impedance control is significant alongside force generation in stable environments [6]. The co-occurrence of force and impedance is a challenge for isolating their control during a neuroimaging experiment.

To first explore how individuals should regulate both force and impedance control in reaching tasks with stable external forces, we use the Stochastic Optimal Open-Loop Control (SOOC) framework [7]. We adapt SOOC to describe our wrist-pointing task and evaluate responses across a range of cost functions and noise parameters. Insights from these simulations are used to introduce and evaluate four force conditions in an experimental 2-DOF wrist-pointing task to maximally decouple force and impedance control. Subsequently, we evaluate the effectiveness of these proposed force conditions to elicit distinct neural activation through a virtual neuroimaging experiment. This combination of neuromuscular modeling, behavioral experiments, and simulated neural activity establishes the foundation for a future neuroimaging study, aiming to precisely locate the neural correlates of force and impedance control of the wrist. The broader impacts of this work are critical in enhancing the classification and diagnosis of neural impairments, while paving the way for more personalized and effective neurorehabilitation treatments.

## II. THEORETICAL FRAMEWORK

To quantify neural function contributing to joint stiffness modulation via co-contraction, one might consider a study design with two types of tasks, such as one with low instability (i.e., a no force task) and one with high instability (i.e., a divergent force task). While these tasks may be sufficient if stiffness is commanded at the joint level, they are inadequate at the muscle level. With these tasks alone, we cannot distinguish whether the ensuing neural signal is attributable to the cortical control of muscle co-contraction (i.e., synergistic co-activation of agonist-antagonist muscle pairs that leads to increased joint impedance) or to separate, independent increases in activation of agonist and antagonist muscles to produce greater forces. Thus, it is not possible with only these two tasks to determine if there are regions in the brain that contribute to the neural control of cocontraction or if muscle-specific regions are responsible for these commands. A constant force task could address this limitation, with the expectation that compensation would primarily result from updates in the internal model based on predicted task dynamics. In an ideal case, the constant force condition would result only in changes in net torque relative to the no force condition, isolating commands specific to agonist force generation (Fig. 1A). However, our prior work [6] cautions against assumptions of negligible impedance control in dynamic wrist movements.

**Fig 1.**
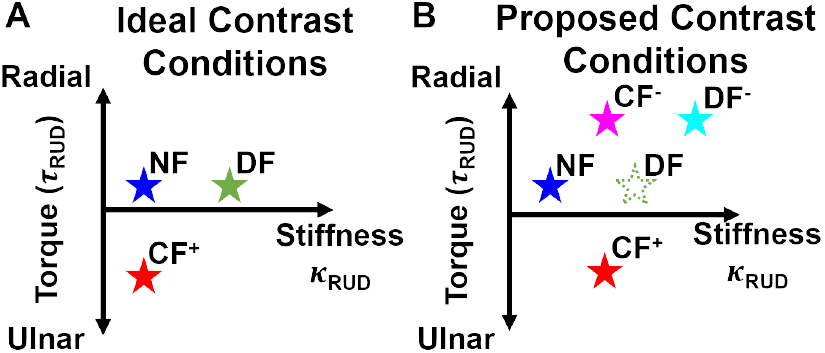
Hypothesized relationship between torque and stiffness during a wrist-pointing task. (A) In the ideal case, the effect of stiffness can be isolated by contrasting the neural activity between DF and NF, and torque can be isolated by contrasting CF and NF. (B) Our experimental task includes the conditions with solid stars.

We first investigate how torque and stiffness may co-vary with the magnitude of applied perturbations using a neuromuscular model and decide what experimental conditions are suitable to maximally dissociate them. This approach enhances our ability to isolate and interpret neural commands associated with force generation and impedance control.

### A. Predictions Using a Neuromuscular Model

The need for impedance control is well documented for unstable or unpredictable tasks, where muscle co-contraction improves task stability and minimizes trajectory errors [2]. However, it remains unclear whether impedance control is also regulated in stable tasks, where the increased metabolic energy costs associated with co-contraction might outweigh the potential benefits introduced by greater execution accuracy. This distinction is important because if impedance control is indeed present in stable tasks, we will need to quantify the behavioral relationship between impedance and torque to define proper neural contrast conditions to identify regions of the brain specific to each process.

We use the Stochastic Optimal Open-Loop Control (SOOC) framework [7] to probe the relationship between torque and stiffness in response to stable perturbations. We opted to use the SOOC framework because it is well suited to account for muscle impedance. SOOC predicts optimal feedforward motor commands by minimizing a cost function that penalizes effort and final variance. In this formulation, the main factors that determine optimal torque and stiffness values are the relative weights on final error and stiffness relative to torque. Motor noise is also added to the system and primarily affects the propagation of the covariance. The methods for using this framework are published in [7]. We modeled a single-joint system, representing the wrist, with two degrees of freedom in flexion-extension (FE) and radial-ulnar deviation (RUD). We considered a wrist extension task with either no externally applied forces (NF: *τ*_*FE*_ = *τ*_*RUD*_ = 0) or with a constant force applied perpendicular to movement direction (CF: *τ*_*RUD*_ = 0.6 Nm).

We found that across various cost functions, the solutions that best minimize effort and variance show an increase in *κ*_*RUD*_ in CF relative to NF (Fig. 2). These results, supported by our previous work [6], suggest that stiffness of the wrist joint is regulated even in the presence of a stable external perturbation. Therefore, at the neural level, it is possible that changes in neural activation between these NF and CF tasks may be explained by groups of neurons responsible for regulating torque, stiffness, or both. This motivates our work to carefully select experimental force conditions that allows us to decouple these processes.

**Fig 2.**
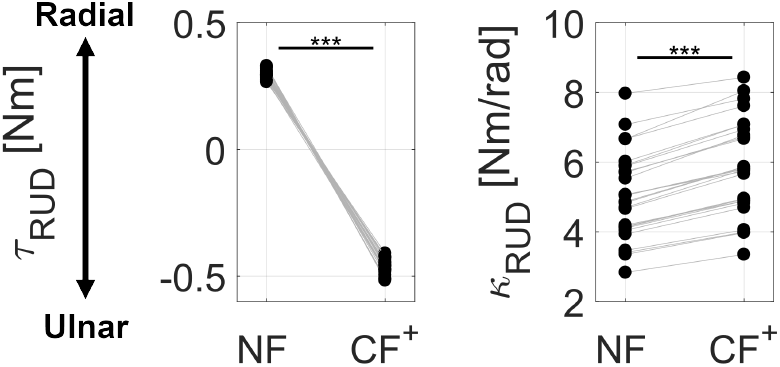
Optimal SOOC-predicted mean torque and stiffness responses. Each gray line/dot represent a single combination of cost function and noise parameters. We tested three levels of cost function weights on final variance (*q*_*f*_ = {0.5, 1, 2}) and stiffness relative to torque (*α* = {0.5, 1, 2 }). Results are shown for signal-dependent noise *d* = {0.2, 0.4, 0.6} and constant noise *σ* = 0.1. *** indicates a difference with *p <* 0.001.

### B. Task Selection

Recall that in an ideal case, the magnitude of the applied force in a CF condition would regulate only torque responses and not stiffness. Qualitatively, this would be indicated by a vertical shift in Fig. 1A. However, our model results indicate that stiffness is also regulated with applied forces (Fig. 2). This would place the CF task slightly to the right of NF on the torque-stiffness plot (Fig. 1B). This distinction is critical because we no longer assume that the CF task results in torque or stiffness values that match those of the divergent force (DF) task. Thus, we need to experimentally determine the conditions that elicit a distinct torque and stiffness response.

In all conditions, forces will be applied via a 2-DOF wrist exoskeleton, the MR-SoftWrist, where interaction forces are controlled through a series elastic actuator [8]. We plan to include the CF condition in two directions (CF^+^ and CF^*−*^) to distinguish the neural commands associated with producing torque in any direction and the commands directed to a specific muscle to produce direction-specific torque. In the CF^+^ condition, force is applied in the positive RUD direction, meaning forces push the hand in the direction opposite to gravity. Conversely, in the CF^*−*^ condition, force is applied in the negative RUD direction, or in the same direction as gravity. We choose the magnitude of the force to be 0.3 Nm, which should be strong enough to yield statistically different results from the NF condition, but not too strong to be overly fatiguing after many trials.

In place of a traditional divergent force condition, we propose a divergent force condition with an offset (DF^*−*^) to act as the high stiffness condition, such that *τ*_*RUD*_ = *kθ*_*RUD*_ +*c*. When *c <* 0, the majority of the workspace is located in a region where the perturbation torque is applied down (Fig. 3), but the instability remains because the magnitude of the force scales with RUD position. This proves advantageous for our study because, with a predictable force direction, we can designate specific muscles as agonists or antagonists, enabling the use of the activation of the pure antagonist muscle [6] as an index of muscle co-contraction. Note that this definition is not possible with a standard divergent force, such as in [2], because the perturbation direction cannot be predicted. We chose *c* = *−* 0.3 Nm, such that in the straight line trajectory between targets (*θ*_*RUD*_ = 0), the force experienced is the same as the CF^*−*^ condition. The position-dependent gain *k* is set as 0.085 Nm/deg, which is strong enough to introduce sufficient instability. Thus, we expect a similar torque response to the CF^*−*^ condition and an increase in stiffness resulting from the stiffness necessary to counter both the offset and the added instability of the position-dependent force field (Fig. 1B).

**Fig 3.**
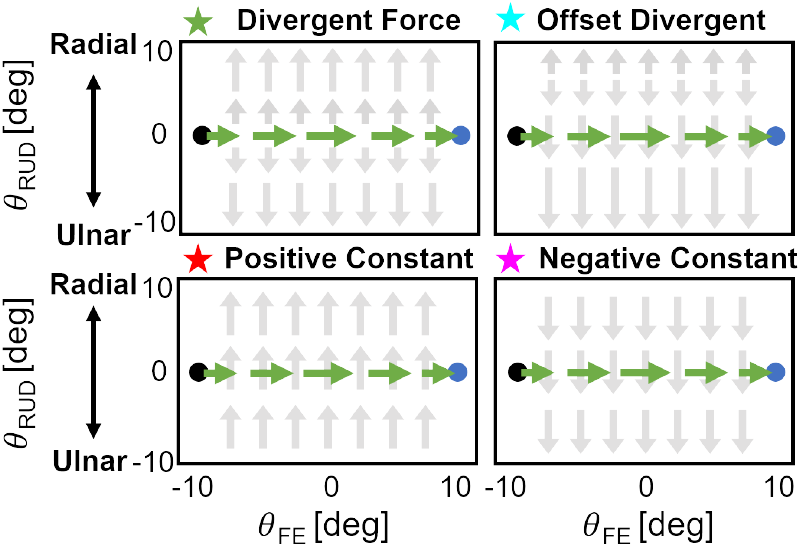
Proposed experimental force fields. In each task, the goal is to move the cursor from the black target to the blue target in a straight line.

## III.BEHAVIORAL EXPERIMENT

### A. Experimental Procedure

Seven healthy young adults (3 male, age: 26 ± 3 years, all right-hand dominant) free from neurological or musculoskeletal injury participated in this single-session study. This study was approved by the Institutional Review Board of the University of Delaware, IRB no. 1936400-2.

The experimental protocol is shown in Fig. 4. At the beginning of the session, EMG sensors were placed on the participant’s four major wrist muscles: flexor carpi radialis (FCR), flexor carpi ulnaris (FCU), extensor carpi ulnaris (ECU), and extensor carpi radialis (ECR). Sensors were placed on the participant’s dominant arm (Fig. 4C-D). We used Delsys Trigno Duo electrodes (sampling frequency: 2148 Hz), and the EMG signal was time-synced to task performance data (sampling frequency: 1000 Hz) via a common analog signal. Electrodes were placed along the muscle belly at the location of the largest agonist muscle contraction. The forearm was prepared for electrode placement by shaving the arm hair and cleaning the skin with alcohol.

**Fig 4.**
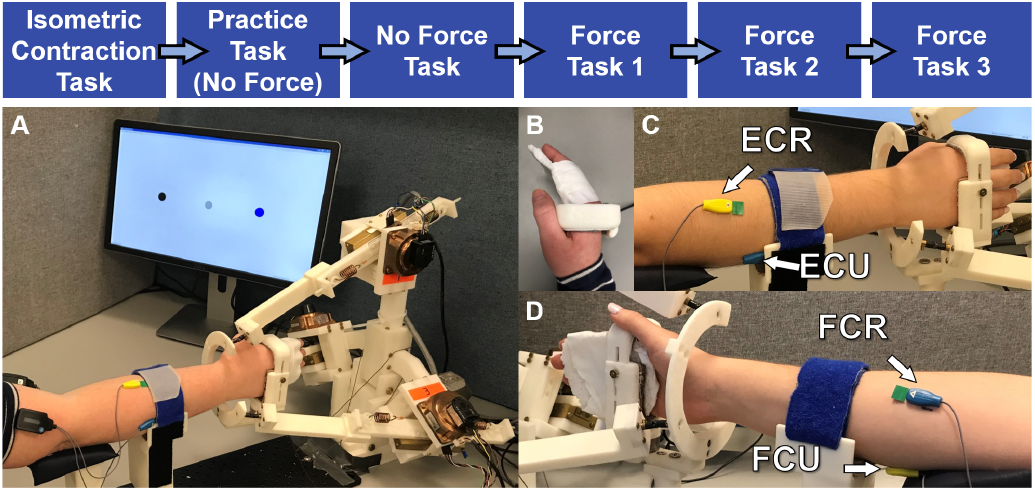
Top: Experimental task sequence; Bottom: (A) MR-SoftWrist during task performance, (B) top view of device handle, where fingers are secured in a flexed position, (C-D) EMG electrodes placed on four wrist muscles.

EMG activation was first measured during isometric contractions. Participants were cued to apply and hold a torque of 1 Nm for 7 seconds three times in each direction (flexion, extension, radial deviation, and ulnar deviation). This procedure was repeated twice: once in a neutral wrist posture (*θ*_*F E*_ = *θ*_*RUD*_ = 0 deg) and again in a flexed posture (*θ*_*F E*_ = 5 deg, *θ*_*RUD*_ = 0 deg). Agonist muscle contractions were used to confirm correct electrode placement, quantify signal quality, and normalize signals across participants.

In all motor tasks, participants used the MR-SoftWrist (Fig. 4A), an MRI-compatible wrist robot [8]. Participants secured their hand in the robot’s handle which clamps around their knuckles without restricting their fingers (Fig. 4B). A small paddle was inserted into the handle to encourage a slightly flexed posture and prevent a power grip [9]. Participants were instructed to focus on performing wrist movements while keeping their fingers as relaxed as possible to minimize coactivation with finger muscles.

Participants controlled a cursor on a monitor by moving the handle in FE and RUD to move the cursor left/right and up/down, respectively. Participants were cued to move the cursor (radius: 1.5 deg) in a straight line between two circular targets (radii: 1.75 deg) located at (±10,0) deg in (FE, RUD). Trial onset was cued by a change in target color from black to blue. Once the participant moved the cursor 1.5 deg laterally toward the target, the cursor was hidden and the applied force was initiated, if applicable. The cursor was hidden to minimize corrections due to online feedback and encourage feedforward motor planning. The trial ended when the participant passed the lateral position of the target (*θ*_*FE*_ = ±11 deg) and the robot assisted the user back to the target via a convergent force field (virtual stiffness = 100 Nm/deg). Two forms of feedback were displayed to the user at the end of each trial: speed feedback and position error feedback. Speed feedback was displayed by turning the target red if the movement duration was greater than 500 ms or green if it was less than 200 ms. Otherwise, the target remained black. Error feedback was displayed via a small yellow circle (radius: 1 deg) located directly above or below the target, at the distance of the maximum RUD error during the trial. Both forms of feedback were displayed for 0.5 s after every trial and followed by a random inter-trial interval selected from a normal distribution *N* (0.75, 0.25) s, bounded between [0.2, 0.8] s before cueing the next trial.

Participants practiced the task in a transparent (no force) condition to familiarize themselves with the task, understand the displayed feedback, and ask any questions. After familiarization, participants completed four blocks of 200 trials, with about five minutes of rest in between blocks to minimize muscle fatigue. We chose 200 trials to account for a sufficient number of trials to reach steady-state conditions (first 100 trials) and have enough steady-state trials to provide a sufficient signal-to-noise ratio (last 100 trials). The first block was always an NF task to measure baseline performance, and the next three blocks were presented in a randomized order and included a positive constant force block (CF^+^), a negative constant force block (CF^*−*^), and an offset divergent force block (DF^*−*^) (Fig. 3, 4), as described in Section II-B. For participant safety and continuous operation, the divergent force condition was saturated at ±1 Nm.

### B. Data Analysis

Signal-to-noise ratio (SNR) was calculated for the isometric contraction task to confirm electrode placement and quantify signal quality. We defined SNR for each muscle *i*:

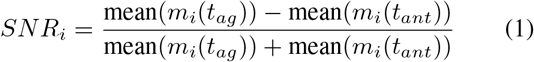

where *m* represents the mean EMG signal during a contraction, and *t*_*ag*_ and *t*_*ant*_ are the times when *i* is agonist or antagonist, respectively. For any muscle, SNR is +1 if the muscle is active only in its agonist direction, 1 if the muscle is only active in its antagonist direction, or 0 if the muscle is equally active in its agonist and antagonist directions.

For trial-by-trial data analysis, trial onset (*t*_0_) was defined as the instant the cursor passed a position threshold of 1.5 deg away from the starting target. Trial end (*t*_*end*_) was defined as the instant the cursor passed a position threshold of 9 deg. To normalize across trials, we defined three key time windows: early, middle, and late trial. The early window starts 150 ms before *t*_0_ and continues until *t*_0_. The middle starts at *t*_0_ and ends when maximum flexion-extension velocity is reached (*t*_*vmax*_), and the late trial starts at *t*_*vmax*_ and ends at *t*_*end*_.

Flexion and extension trials were considered separately, and out of each block of 200 trials, we considered only the last 100 trials to remove the transients of learning each force condition. EMG signals are bandpass filtered at 30-500 Hz, rectified, and the envelope is taken via a 10 Hz low-pass, zero-shift 4th order Butterworth filter.

1) Muscle Level: As an index of co-contraction, we considered the activity of the pure antagonist [6], that is, the muscle that is antagonistic in both movement direction and perturbation direction. During the NF task, gravity acts to pull the hand downward. In this case, to meet the task goal of maintaining a neutral RUD posture at the target,the radial deviator muscles act as agonists pulling the hand upwards, making the ulnar deviators the antagonists. Our task conditions were carefully defined such that the applied force is also directed downward in both the CF^*−*^ and DF^*−*^ conditions, so the ulnar deviators also act as antagonists in these tasks. Thus, in these tasks, the pure antagonist is ECU during flexion trials and FCU during extension trials.
2) Joint Level: We used the isometric contraction task to estimate muscle-specific scaling factors to compute muscle forces and joint torque and stiffness via a forward dynamics estimation procedure [10]. For each posture tested, we obtained muscle-specific moment arms from OpenSim [11]. The moment arms were compiled into a Jacobian matrix ***J***, where the rows *i* correspond to a wrist degree of freedom (FE or RUD) and the columns *j* correspond to a specific muscle, resulting in a 2 *×* 4 matrix. Then, EMG measurements *m* were related to joint torque:

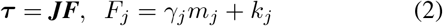

Muscle scaling and offset factors (***γ*** and ***k***, respectively) were determined by relating measured torques from the F/T sensor to these estimates with a least-squares regression:

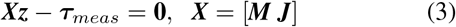

where ***z*** = [***γ***; ***k***] is a 8 *×* 1 vector and ***M*** is a 2 × 4 matrix with elements *M*_*ij*_ = *J*_*ij*_*m*_*j*_.

Joint stiffness was estimated using the short-range stiffness approximation [12]:

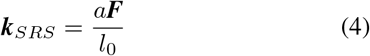

where *a* is a dimensionless scaling constant (*a* = 23.4) [12] and *l*_0_ is the optimal muscle fiber l ength a t maximum activation, as defined i n O penSim [11]. J oint l evel stiffness was computed as:

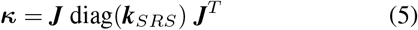

We also estimated the resulting contributions to torque in the direction of perturbation at the joint level as:

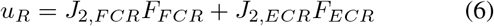

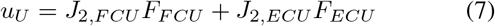

where *J*_2,*j*_ is the muscle’s RUD moment arm and *F*_*j*_ is the estimated force. These quantities were evaluated to define a compound metric of agonist and antagonist activation, useful for defining co-contraction at the joint level.

### C. Statistical Testing

We used the Wilcoxon Signed Rank test to evaluate torque and stiffness across the different conditions in the early time window, selected to focus on planned actions rather than online feedback responses. At the muscle level, we used the mean activations of the pure antagonist as a proxy for joint stiffness. At the joint level, we estimated mean *κ*_*RUD*_, *τ*_*RUD*_, *u*_*R*_, and *u*_*U*_. To focus on changes induced by each force condition and remove subject-specific biases, the mean in each force condition was subtracted by the mean in NF.

### D. Results

SNR was calculated separately when the muscle is agonist in the FE and RUD directions during the isometric contraction task (1). In the FE direction, the mean and standard deviation of the SNR of each muscle across all seven participants was: FCR: 0.81 ± 0.16, FCU: 0.58 ± 0.16, ECU: 0.78 *±* 0.16, and ECR: 0.83 *±* 0.15. In the RUD direction, SNR of each muscle was: FCR: 0.10 *±* 0.42, FCU: 0.83 *±* 0.21, ECU: 0.68 *±* 0.11, and ECR: 0.74 *±* 0.16.

In flexion trials, where ECU is the pure antagonist in NF, CF^*−*^, and DF^*−*^, we observe an increase in its mean activity in CF^*−*^ and DF compared to NF in all participants. This result is significant at the group-level (CF^*−*^-NF: *p* = 0.0078, DF-CF: *p* = 0.0078). The pure antagonist activity (ECU) is shown for three participants in Fig. 5. In extension trials, where FCU is the main pure antagonist, we observe increases in its mean activity from NF to CF^*−*^ in 6/7 participants (not significant at the group level at *α*=0.05: *p* = 0.0547). We also observe significantly greater pure antagonist activity in DF^*−*^ compared to CF^*−*^ in 6/7 participants (*p* = 0.039).

**Fig 5.**
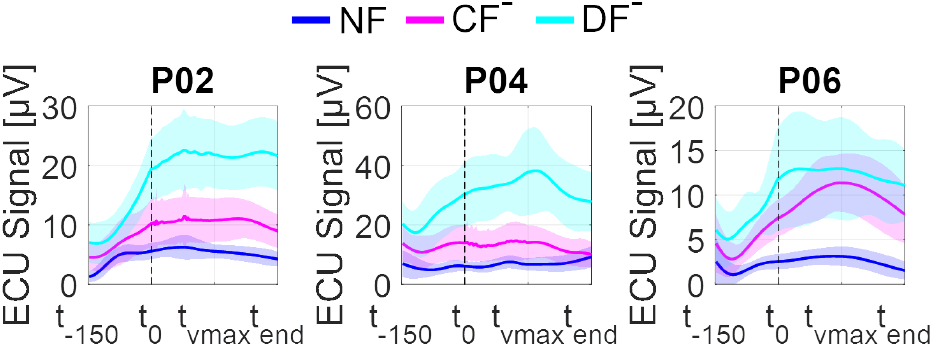
Subject-level pure antagonist responses of ECU for steady-state, flexion trials for three representative subjects. The bold line represents the mean muscle activity, and the shaded area has height of one standard deviation.

At the group-level, we find a significant increase in *k*_*RUD*_ in the normalized DF^*−*^ condition compared to both CF^+^ (*p* = 0.0391) and CF^*−*^ conditions (*p* = 0.0156) in flexion trials (Fig. 6). There are no significant differences in group-level *κ*_*RUD*_ in extension trials, but there is a trend of increasing *κ*_*RUD*_ from CF^*−*^ to DF^*−*^ in 6/7 participants.

**Fig 6.**
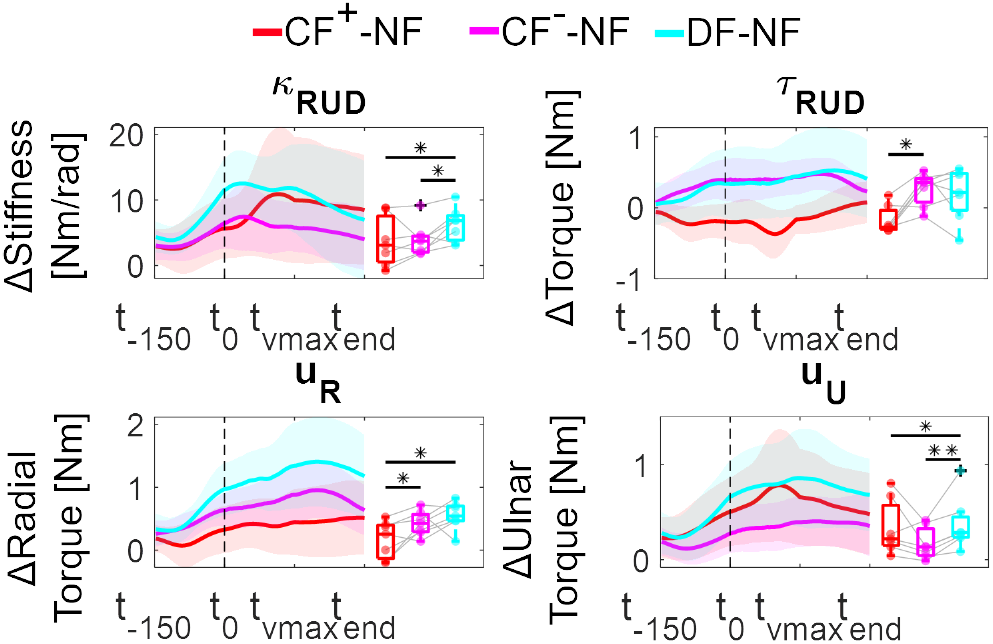
Group-level torque and stiffness estimates for steady state, flexion trials, normalized by NF activity. The boxplots show the distribution of each subject’s mean activity in the early time window (*t*_*−*150_ : *t*_0_). *∗* indicates *p <* 0.05, *∗∗* indicates *p <* 0.01.

**Fig 7.**
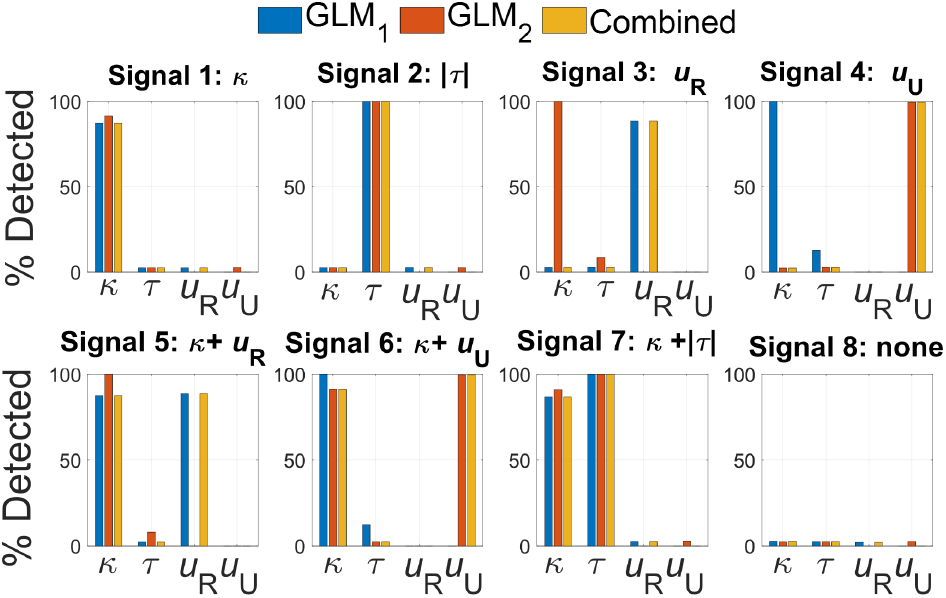
Detection rate of each state when SNR=1 and N=150.

## IV. SIMULATED NEUROIMAGING EXPERIMENT

The ultimate goal of this project is to characterize patterns of torque and stiffness expression such that their neural commands can be maximally decoupled and identified during functional magnetic resonance imaging (fMRI). Here, we use a general linear model approach [13] to analyze a simulated fMRI experiment to examine if our proposed force conditions are sufficient to distinguish between the neural representations of stiffness, torque in any direction, and direction-specific torque. To do this, we simulate possible neural signals that might produce the torque and stiffness patterns measured experimentally and evaluate if we can accurately identify activation associated with each state.

We intend to use a block design, where we will consider all steady-state trials from each condition. We expect the hemodynamic response to remain high for the duration of the block because the blood-oxygen-level-dependent (BOLD) signal that is measured during fMRI is slow to respond (temporal resolution: *∼*1-3 seconds) and thus will not have time to decrease between trials. We define eight possible voxel types, which respond to *κ*, |*τ*|, *u*_*R*_, *u*_*U*_, *κ*+*u*_*R*_, *κ*+*u*_*U*_, *κ* + |*τ*|, or none of these. Then, we predict how these voxels will respond to each experimental condition based on the group-level experimental responses in the early time window during flexion trials. For each of the simulated voxels, we added zero-mean Gaussian noise with a standard deviation equal to the maximum expected signal for each regressor (SNR=1). Each simulation was run 1,000 times to evaluate how well each model correctly identified each voxel type.

We defined General Linear Models (GLMs) for regression onto our simulated fMRI data. Our regressors of interest are stiffness (*κ*_*RUD*_), absolute torque (|*τ*_*RUD*_|), and direction-specific torque (*u*_*R*_ and *u*_*U*_). Here, *u*_*R*_ is the contribution to torque in the radial direction (6), and *u*_*U*_ is the contribution in the ulnar direction (7). The linear dependency between explanatory variables makes the inclusion of all four regressors result in a reduced-rank experimental matrix, so we define two GLMs to fully explain all voxel types:

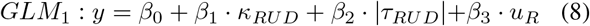

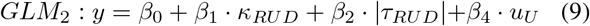

### A. Data Analysis

We performed a regression with the simulated neural activity and each GLM to calculate the estimated contribution of each regressor (*κ*_*RUD*_, |*τ*_*RUD*_|, *u*_*R*_, *u*_*U*_) to the measured signal, known as the *β* estimates. We considered the bounds on the confidence interval for each *β* estimate to determine if each regressor is positively or negatively correlated with the simulated neural activity. We calculated the detection rate by considering the number of cases where *β >* 0 for the correct voxel types out of the total number of simulations run.

In our approach, no single model can directly explain the signal from all the considered voxel types, due to the reduced-rank problem described earlier. When the explanatory variable for a specific voxel is not directly included in the model, a negative *β* coefficient is typically estimated,so to effectively combine the information conveyed by the two GLMs into one model, we started with GLM_1_ (8) and examined the *β* weights estimated for each regressor. If *β <* 0, then we used the estimated *β* weights from GLM_2_ (9) for that signal. This combined signal is useful for fMRI analysis, where the true voxel types are not known.

### B. Results

When N = 100 trials, we find that across all voxel types, there is a minimum successful detection rate of 70.1% and a maximum false positive detection rate of 2.7%. The success rate improves with the length of each block (N=150: 86.69%, N=200: 94.27%), while the false positive rate remains low (N=150: 2.8%, N=200: 2.69%).

## V. DISCUSSION

This study outlines steps in designing a neuroimaging experiment to dissociate the neural substrates of torque and stiffness. We designed experimental task conditions to elicit a unique combination of torque and stiffness, based on insights from prior work and neuromuscular modeling. Our experimental results reveal unique muscle responses in flexion trials (Fig. 6), while in extension trials, *κ*_*RUD*_ showed no significant difference across force conditions. This aligns with prior work that highlights stronger co-contraction effects in wrist flexion movements [14]. This suggests that these force conditions may only yield significant differences in co-contraction in flexion trials, so we will only include flexion trials in our fMRI study and command the robot to return to the starting position after every trial.

The DF^*−*^ condition, unique to this study, proves useful in cases where EMG measurements are noisy. Our primary outcomes of torque and stiffness are estimated via EMG at the joint level, which is susceptible to errors due to variations in normalization and noise levels across EMG electrodes. Defining DF^*−*^ with the offset means that the force will pull downward in the majority of the workspace, enabling the identification of specific muscles as pure antagonists. This approach allows us to contrast the effects of force conditions within a single muscle, providing an additional metric of co-contraction that is not influenced by normalization errors. While we observed that FCR had a low SNR on average in the RUD direction (0.10 ± 0.42), FCR was only the pure antagonist muscle in CF^+^, so we can still fully evaluate co-contraction via the ranked order of activity of the pure antagonist in NF, CF^*−*^, and DF^*−*^. We observe the expected ranking of pure antagonist activity in every participant in flexion trials and 6/7 participants in extension trials, in agreement with our joint level estimation of *κ*_*RUD*_ (Fig. 6).

Next, we computed simulations of neural activity and assessed whether these responses could be isolated via GLMs. Our simulations indicate that at least 150 steady-state trials are needed to obtain results powered to detect all voxel types at the 80% level, a standard threshold for power detection. In conclusion, we show that the NF, CF^+^, CF^*−*^, and DF^*−*^ conditions evoke distinct muscle responses and are well suited for study during fMRI. This represents an important first step in isolating the neural origins of co-contraction, which has numerous implications for better understanding motor control processes and their impact on motor pathways.

## REFERENCES

[1] D. M. Wolpert, Z. Ghahramani, and M. I. Jordan, “An Internal Model for Sensorimotor Integration,” Science, vol. 269, pp. 1880–1882, 1995.

[2] E. Burdet, R. Osu, D. W. Franklin, T. E. Milner, and M. Kawato, “The central nervous system stabilizes unstable dynamics by learning optimal impedance,” Nature, vol. 414, pp. 446–449, 11 2001.

[3] R. Osu, E. Burdet, D. W. Franklin, T. E. Milner, and M. Kawato, “Different Mechanisms Involved in Adaptation to Stable and Unstable Dynamics,” J Neurophysiol, vol. 90, pp. 3255–3269, 11 2003.

[4] M. L. Latash, “Muscle coactivation: definitions, mechanisms, and functions,” J Neurophysiol, vol. 120, pp. 88–104, 3 2018.

[5] W. Sheng, S. Li, J. Zhao, Y. Wang, Z. Luo, W. L. A. Lo, M. Ding, C. Wang, and L. Li, “Upper Limbs Muscle Co-contraction Changes Correlated With the Impairment of the Corticospinal Tract in Stroke Survivors: Preliminary Evidence From Electromyography and Motor-Evoked Potential,” Frontiers in Neuroscience, vol. 16, 6 2022.

[6] A. J. Farrens, K. Schmidt, H. Cohen, and F. Sergi, “Concurrent Contribution of Co-Contraction to Error Reduction During Dynamic Adaptation of the Wrist,” IEEE Transactions on Neural Systems and Rehabilitation Engineering, vol. 31, pp. 1287–1296, 2023.

[7] B. Berret and F. Jean, “Stochastic optimal open-loop control as a theory of force and impedance planning via muscle co-contraction,” PLoS Computational Biology, vol. 16, 2 2020.

[8] A. Erwin, M. K. O’Malley, D. Ress, and F. Sergi, “Kinesthetic Feedback during 2DOF Wrist Movements via a Novel MR-Compatible Robot,” IEEE Transactions on Neural Systems and Rehabilitation Engineering, vol. 25, pp. 1489–1499, 9 2017.

[9] W. L. Popp, O. Lambercy, C. Müller, and R. Gassert, “Effect of Handle Design on Movement Dynamics and Muscle Co-activation in a Wrist Flexion Task,” Int J Ind Ergonom, vol. 56, pp. 170–180, 11 2016.

[10] D. R. Smith, C. A. Helm, A. Zonnino, D. J. McGarry, C. L. Johnson, and F. Sergi, “Individual Muscle Force Estimation in the Human Forearm using Multi-Muscle MR Elastography (MM-MRE),” IEEE T Bio-Med Eng, vol. 70, pp. 3206–3215, 11 2023.

[11] K. R. Saul, X. Hu, C. M. Goehler, M. E. Vidt, M. Daly, A. Velisar, and W. M. Murray, “Benchmarking of dynamic simulation predictions in two software platforms using an upper limb musculoskeletal model,” Computer methods in biomechanics and biomedical engineering, vol. 18, p. 1445, 10 2015.

[12] X. Hu, W. M. Murray, and E. J. Perreault, “Muscle short-range stiffness can be used to estimate the endpoint stiffness of the human arm,” J Neurophysiol, vol. 105, pp. 1633–1641, 2 2011.

[13] J. B. Poline and M. Brett, “The general linear model and fmri: Does love last forever?,” NeuroImage, vol. 62, pp. 871–880, 8 2012.

[14] G. N. Forman, D. A. Forman, E. J. Avila-Mireles, J. Zenzeri, and M. W. Holmes, “Investigating the Muscular and Kinematic Responses to Sudden Wrist Perturbations During a Dynamic Tracking Task,” Scientific Reports, vol. 10, 12 2020.

